# Time-resolved, integrated multi-omic analysis reveals central role of amino acid pathways for defense responses in *Arabidopsis thaliana*

**DOI:** 10.1101/2024.08.27.609849

**Authors:** Charlotte Joller, Klaus Schläppi, Joelle Sasse

## Abstract

Plants respond to biotic stresses by altering gene expression and metabolism. However, how fast different tissues respond to microbial presence, and how similar these responses are is mostly unresolved. Here, we treat *Arabidopsis thaliana* with elicitors and investigate time-resolved changes in shoot, root, and root-derived (exudate) metabolite profiles. We find that root responses precede shoots and that first metabolite changes take place after 1.5 h and persist for 3 d. Exudates respond within 4 h (earliest timepoint available) to elicitor presence. This response diminishes when plants are pulse-treated but persists for continuously treated plants. Defense compounds such as phenylpropanoids increase after 1.5-4 h. Amino acids were iden6fied as central players in defense: they increase after 1.5 h in shoots, roots, and exudates. Transcriptome analysis at 4 h and 1 d and integrated, multi-omic analysis of transcription and metabolome suggest that aromatic and aliphatic amino acids are central players in defense responses. As their transcriptional and metabolite increases are fast and persisting over days for most amino acids, we propose amino acids as early indicators for biotic stress monitoring.

## Introduction

Plants are sessile organisms that alter metabolism in response to pathogens, as they cannot evade the stress. Metabolic responses to a biotic stimulus stress are microbe and host specific^1^. A hallmark of defense responses against a pathogen is the production of specialized compounds, which are broadly classified as isoprenoids (e.g. terpenes), shikimates (e.g. phenylpropanoids), alkaloids, and glucosinolates, depending on their pathway of origin^1,2^. Isoprenoids are synthesized from C5-units originating from Acetyl-CoA, a product of central carbon metabolism^1^. Phenylpropanoids and alkaloids derive from aromatic amino acids^1^, and glucosinolates similarly from aromatic or aliphatic amino acids^2^. Thus, when a pathogen is encountered, primary and secondary metabolite levels are changed. Cultivars or ecotypes resistant against a pathogen typically feature a metabolite profile distinct from their susceptible counterparts^3–5^: resistant cultivars show increased levels of defense compounds such as phenolics, flavonoids, and salicylic acid^3^, and elevated carbohydrate, amino acid, and carboxylic acid levels^6^. When investigating metabolite changes in a host infected with a pathogen, a major issue is the presence of the pathogen with its own metabolism generating and turning over compounds. Thus, many studies mimic pathogen presence with elicitors, which are small molecules stimulating plant defense. *Arabidopsis thaliana* treated with the elicitors flg22, chitosan, or a combination thereof showed changes in phenylpropanoids, glucosinolates, terpenoids, alkaloids, and phenolic compounds^7^. Arabidopsis callus tissue similarly responded to lipopolysaccharide treatment, another elicitor, with changes in glucosinolates, alkaloids, and cinnamic acid^8^, and tomato leaves infiltrated with flg22 and flgII-28 found changes in flavonoids, glycoalkaloids, lipids, organic and amino acids^9^.

It is to be expected that these metabolite changes are preceded by transcriptional and proteomic changes. Indeed, plants treated with elicitors alter their transcriptional profile, as shown for example for Arabidopsis seedlings treated with seven different elicitors^10^. Early transcriptional responses mostly target defense-related genes, whereas later responses encompass changes of growth and development^10,11^. A major question in experiments investigating gene expression and metabolite changes in response to a stimulus is the timing of the response: For pathogen-treated plants, metabolomic samples are often collected one or few days after treatment^3,6^, likely because the pathogen needs time to enter the host tissue, establish in the tissue, and elicit a response strong enough to be measured on a whole-tissue level. In elicitor-treated plants, tissues are also collected in the same timespan^7–9^. However, when transcriptional responses are investigated, samples are collected minutes to hours after treatment, and generally not after days: In a timecourse experiment performed with Arabidopsis seedlings treated with elicitors, samples were collected from 5-180 min, and strongest response was detected at 30-180 min^10^. Although transcriptional responses logically precede metabolite responses, it is to date unresolved how quickly these metabolite changes are detected after elicitor treatment, and if various metabolite responses succeed each other, as observed in transcriptional data.

Another crucial factor to consider when investigating plant-pathogen interactions is the tissue investigated. Historically, most research was performed on aboveground tissues, typically on leaves. However, there are a plethora of root pathogens^12,13^. Similar to beneficial and neutral microbes, pathogens are attracted by plant-derived compounds that are exported from roots, so-called root exudates^14^. When challenged with a pathogen, shoots, roots, and exudates metabolite profiles change distinctly^15,16^. Barley challenged with *Fusarium* increased exudation of organic acids^17^, and Arabidopsis responded to pathogen presence with camalexin exudation^18^. The timing of these experiments was comparable to the ones mentioned above, with exudates collected days after infection.

Here, we systematically investigated metabolite responses in shoots, roots, and exudates to elicitors as a proxy of pathogen presence, allowing us to compare responses across sample types. We further collected samples in a timecourse with earliest samples collected 10 min and latest 3 d after treatment to elucidate whether metabolite responses are distinct between early and late timepoints. Last, we coupled metabolite with transcriptional profiles and performed integrated multi-omic analysis. Aside from the expected changes in defense compounds, we highlight the importance of amino acid metabolism in defense.

## Results

### Complex metabolic response of tissues to elicitor treatment

In a first experiment, time-dependency of metabolic changes in shoots and roots in response to elicitors were investigated. Roots of 3-week-old, plate-grown *Arabidopsis thaliana* Col-0 plants were treated with a mixture of 1 µM flg22, 1 µM elf18, and 5 µM pep1 (“fep”, Figure 1A). After the indicated incubation time (0, 10 min, 30 min, 90 min, 4 h, 8 h, 1 d, 2 d, 3 d), roots and shoots were separated, flash frozen, and metabolite profiles were analyzed. Elicitor-treated plants showed a shoot growth retardation as expected (Figure S1A). Of the features detected, 480 compounds were iden6fied via database searches. The tissues comprised of 26% lipids, 14% organoheterocyclic compounds, 12% amino acids, peptides, and analogues, 9% benzenoids, 9% organic acids and derivatives, 8% phenylpropanoids and polyketides, and 5% carbohydrates and conjugates. Below 5% of compounds were assigned to 10 other classes, among them nucleosides, alkaloids, and lignans (Classyfire Superclasses, with the addition of the amino acids Subclass belonging to the organic acids superclass, Figure 1B, Figure S2A). The chemical composition of shoots, roots, mock and fep-treated samples were comparable at this chemical class level (Figure S2A).

**Figure 1.**
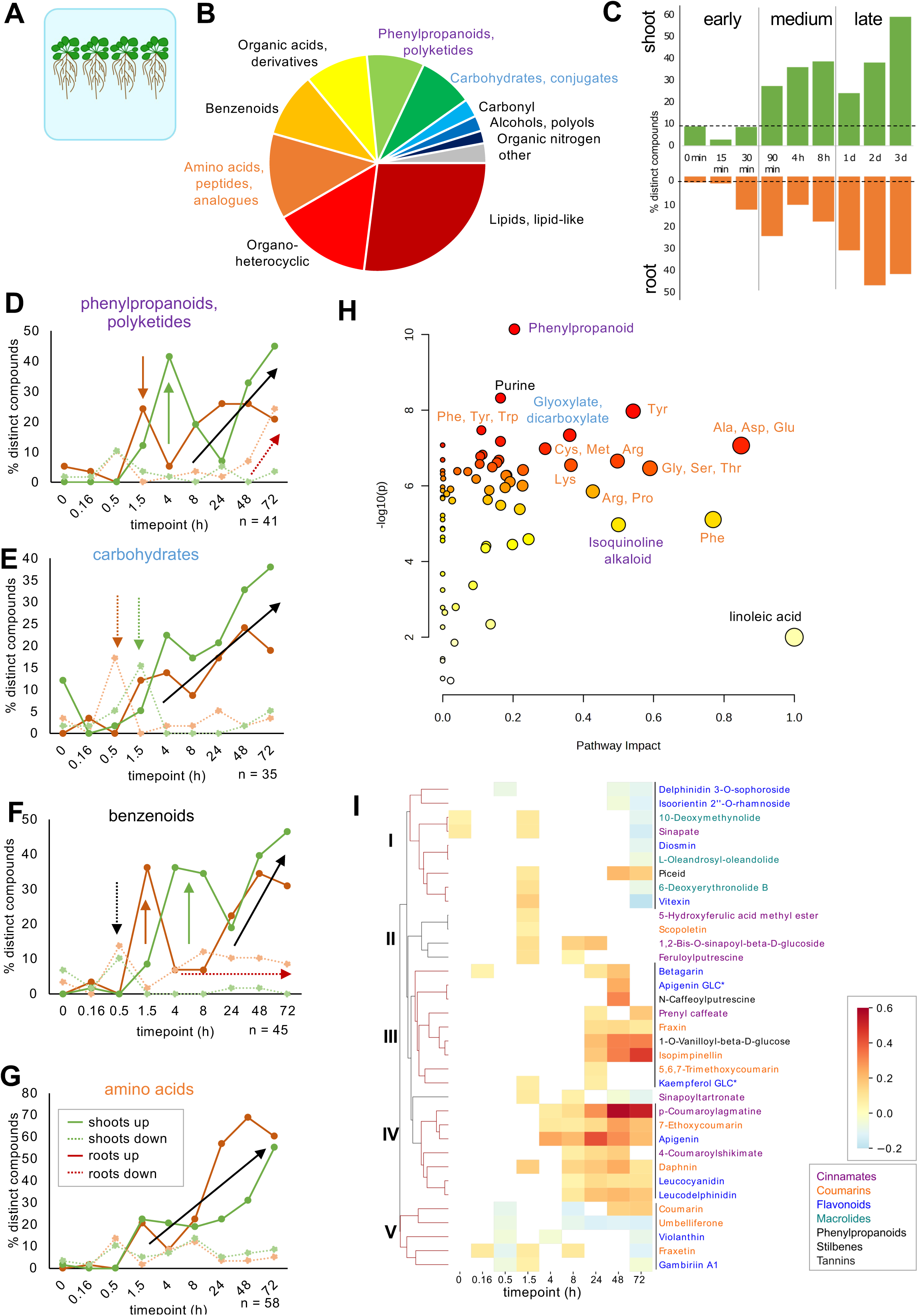
Complex metabolic response in tissues to elicitors. A, Roots of 3-week-old A. thaliana Col-0 were grown on plate, and treated with mock or fep (1 uM flg22, elf18, 5 uM pep1) and sampled after indicated time. B, distribution of chemical classes (Classyfire super/subclass information) in roots. Other: lignans& related, Nucleosides, derivatives& analogues, organosulfur and –metallic compounds, Acetylides, Alkaloids& derivatives, organic 1,3-dipolar comounds. Class distributions are similar in shoots and for compounds distinct between fep and mock in both tissues (Figure S2). Classes depicted in D-G are indicated in bold. Number of metabolites: 480. C, % distinct compounds in fep vs mock treated plants for each timepoints in shoots and roots (Anova/Tukey significance test, p<0.05, n=479-481 metabolites per timepoint and tissue). The dotted line indicates % distinct componds observed at timepoint 0, which served as a background measure. D-G, percent of compounds significantly more or less abundant in fep vs mock-treated tissues (green: shoot, brown: root, solid: more, dashed: less abundant). Number of metabolites is indicated for each class. Arrows highlight patterns observed (same color, pattern as for lines, black: observed in both). N: number of metabolites present in each chemical class. Additional classes are shown in Figure S2. H, pathways changing in roots fep/mock data. Most significant pathways are labeled (amino acid-related pathways highlighted in blue). Additional root pathways are labeled in Figure S3A, shoot pathways are depicted in Figure S3B. I, Clustered heatmap of phenylpropanoids changing significantly in roots over time as fep/ mock treated, averaged by timepoint, log transformed and centered around 0 (red: higher, blue: lower abundant). Color according to Classyfire Class: purple: cinnamates, orange: coumarins, blue: flavonoids, turquoise: macrolides. Others: phenylpropanoids, stilbenes, tannins. Additional maps are shown in Figure S5.

Up to 10% of compounds were changing in the early timepoints 0-30 min, likely representing background, resulting in an overlap of these timepoints in a principal component analysis (PCA, Figure S1B). In medium timepoints 90 min – 8 h, up to 25% of root and 40% of shoot compounds changed significantly between mock and elicitor-treated plants (Figure 1C, Figure S2A, Anova/Tukey, p-value < 0.5). On a PCA plot, middle timepoints started to separate (Figure S1B). The number of distinct compounds increased to 45% in roots and up to 60% in shoots in late timepoints1 d – 3 d, clearly separating mock and elicitor-treated tissues in PCA plots (Figure 1C, Figure S1B). The number of compounds changing significantly between mock and elicitor-treated tissues increased somewhat linearly over time (Figure 1C). To summarize: tissues responded within minutes-hours to treatment with an elicitor mix. Metabolic changes increased over time and persisted until the end of the experiment at 3 d.

When investigating temporal dynamics of single compound classes, various patterns are observed. For example, 10% of phenylpropanoids decreased in roots and shoots at 30 min, followed by 25% of compounds increasing in abundance first in roots at 90 min, and by 40% of compounds in shoots at 4 h (Figure 1D). Carbohydrates decreased in roots and shoots at 30 min and 90 min, respectively (Figure 1E). Benzenoids similarly decreased at 30 min in both tissues, followed by an increase first in roots, then in shoots (Figure 1F). Amino acids (Figure 1G) as well as most other classes increased at later timepoints (Figure 1D-G, Figure S2B). Overall, a rather small decrease in abundance was observed in many chemical classes at early timepoints, whereas the abundance of many compounds increased at later timepoints. Interestingly, responses in roots often preceded shoots, which might be explained by the elicitor application to roots.

When fold changes fep/mock of root metabolite profiles were analyzed with the MetaboAnalyst Pathway tool, approximately 50% of compounds were matched to a pathway (Table S1). The phenylpropanoid pathway scored with the highest significance, followed by purine and glyoxylate/diarboxylate pathways, and many amino acid-related pathways (Figure 1H, Figure S3A). Carbohydrate-related and other defense-related pathways scored with lower significance (Figure S3A). Shoot metabolite profiles changed similarly to root profiles, also highlighting the importance of amino acids (Figure S3B). However, phenylpropanoids exhibited lower significance levels in shoots compared with roots, whereas glycerolipid and pantotheate/CoA pathways exhibited higher significance (Figure S3B). Mapping the most distinct root pathways onto a KEGG pathway map, it is evident that the different significantly distinct amino acids are well-connected via their respective pathway intermediates (Figure S4A). Many amino acids increase in abundance over time, as do many other key compounds of these pathways (Figure S5B, C). The increase observed for amino acids was more prominent in roots compared with shoots (Figure S5B, C).

Phenylpropanoids are poorly represented on KEGG maps (Figure S4A). They cluster partially with purine metabolism, whereas glyoxylate and linoleic acid metabolism are not connected (Figure S4A). Phenylpropanoid abundance also increases with time (Figure 1I, Figure S5A). Phenylpropanoids comprise many defense compounds, which are expected to change in response to elicitor treatment. We thus asked whether there were distinct responses between different types of phenylpropanoids. For this, phenylpropanoids were grouped into cinnamates, coumarins, flavonoids, and four other classes (Classyfire Class) and the fold change of fep/mock was displayed over time, clustered by similarity in response (Figure 1I). Root phenylpropanoids grouped into five clusters, without any obvious grouping according to class. Many compounds were upregulated at later timepoints (cluster III: 1 d and later, cluster IV: 8 h and later), whereas other compounds were downregulated (cluster I: 3 d, cluster V: across timepoints). In shoots, phenylpropanoids exhibited less prominent changes, resulting in less distinct clustering of compounds (Figure S5A). Shoot phenylpropanoids also increased in abundance over time, and also, no clustering according to Class was observed (Figure S5A).

We detected metabolite changes upon elicitor treatment within few hours. These responses persisted for several days. Root responses were more prominent and preceded shoot responses. A general increase in metabolite abundance with time was observed for many chemical classes, although the relative chemical class distribution did not change with elicitor treatment.

### Transcriptional data supports importance of amino acid metabolism

To compare transcriptional with metabolic responses, plants were grown in a semi-hydroponic setup^19^ and treated at three weeks after germination with mock or elicitors for 4 h or 1 d, representing an early and late timepoint when comparing to the metabolite timecourse. Roots and shoots were pooled, flash frozen, RNA extracted and sequenced. Of 33’610 genes, 21’137 were expressed (63%), and 8249 genes (25% total, 39% of expressed) changed abundance significantly between mock and elicitor-treated plants at either timepoint (Table S2). All treatments clearly clustered apart on a PCA plot (Figure S6). At 4 h, 5961 genes were expressed distinctly (28% of expressed, 49% upregulated), which decreased to 4587 distinctly expressed genes at 1 d (22% of expressed, 52% upregulated, Figure 2A). When clustering significantly changing genes, six clusters were observed: Many genes were upregulated in elicitor-treated plants at 4 h (cluster III), some at 1 d (cluster VI), some at both timepoints (cluster II). Also, many genes were downregulated at 4 h (cluster V) or at both timepoints (cluster IV).

**Figure 2:**
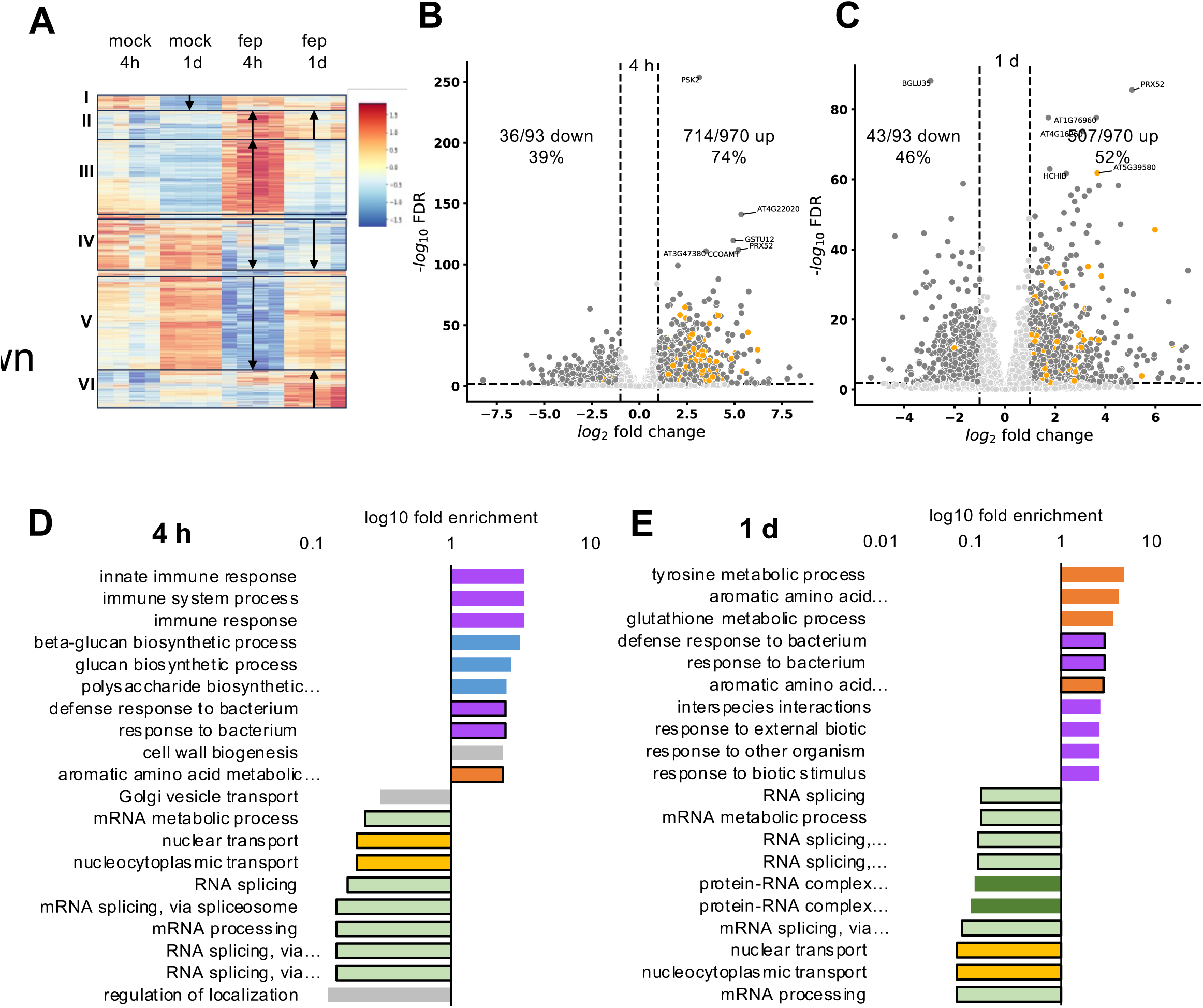
Differential transcription of immune and amino acid genes. RNAseq analysis of 3-week-old pooled roots and shoots treated for 2 h with mock or fep, harvested 4 h or 1 d after the start of the treatment. A, Clustered heatmap of 8259 differentially regulated genes. red: upregulated, blue: downregulated. N = 4 biological replicates. Roman numerals and black boxes indicate cluster number, arrows indicate up- or downregulation of genes in respective treatments. B,C, volcano plots of genes expressed at 4 h (B) and 1 d (C) in grey. Genes present in the 1063 differentially regulated genes iden6fied in Bjornson 2021 are indicated in orange. The number of iden6fied genes up- or downregulated compared to the Bjornson list is indicated on the plot. D, E Top10 differentially up- and downregulated GO terms at 4 h (D) and 1 d (E). GO terms were iden6fied as biological process with Panther, and are colored according to function: green: RNA-related, yellow: nuclear transport, light blue: immunity, dark blue: biotic interactions, purple: carbohydrate metabolism, orange: amino acid, grey: other. Black boxes: GO term iden6fied in both timepoints.

We compared our RNAseq data to an RNAseq experiment of Arabidopsis seedlings treated with 7 distinct elicitors, harvested at 5-180 min, in which 1063 genes were iden6fied that comprise a core group of genes responding to elicitors (Bjornson 2021 Nat Plants). In the 4 h timepoint, 71% of these genes were expressed, with more coverage of up-than down-regulated genes (74% vs 39%, Figure 2B). At 1 d, the number of iden6fied genes dropped to 52%, with similar coverage of up- and downregulated genes (52% vs 46% Figure 2C). Interestingly, the genes most strongly changing in our conditions were not ones iden6fied in this candidate list, which might be due to the different experimental setup.

GO-term enrichment analysis for biological processes in our dataset revealed downregulation of RNA- and nuclear transport-related processes in both timepoints (Figure 2D, E). At 4 h, immune response and defense response terms were enriched as well as glucan and polysaccharide synthesis related genes. At 1 d, some defense-related changes persisted, with other organism/biotic response terms appearing. Interestingly, aromatic amino acid and tyrosine metabolism was detected four times overall across both timepoints.

### Integrated analysis highlights importance of metabolic data

Integrated analysis of metabolomic and transcriptomic was performed with two procedures and a total of four datasets (4 h transcriptomics vs. 4 h roots and shoots, 1 d transcriptomics vs. 1 d roots and shoots). In the first analysis, MetaboAnalyst pathway matching resulted in iden6fication of a set of pathways significantly changing that was similar across all four datasets. Three defense pathways (phenylpropanoids, alkaloids), 2-4 carbohydrate pathways (starch& sucrose, pentose& glucuronate, at 4 h in addition glyoxylate& dicarboxylate and Carbon fixation), and 4-5 amino-acid related pathways (Phe, Tyr, Trp, glutathione, at 1 d in addition Cys& Met) were rated most significant (Figure 3a, Figure S9).

**Figure 3:**
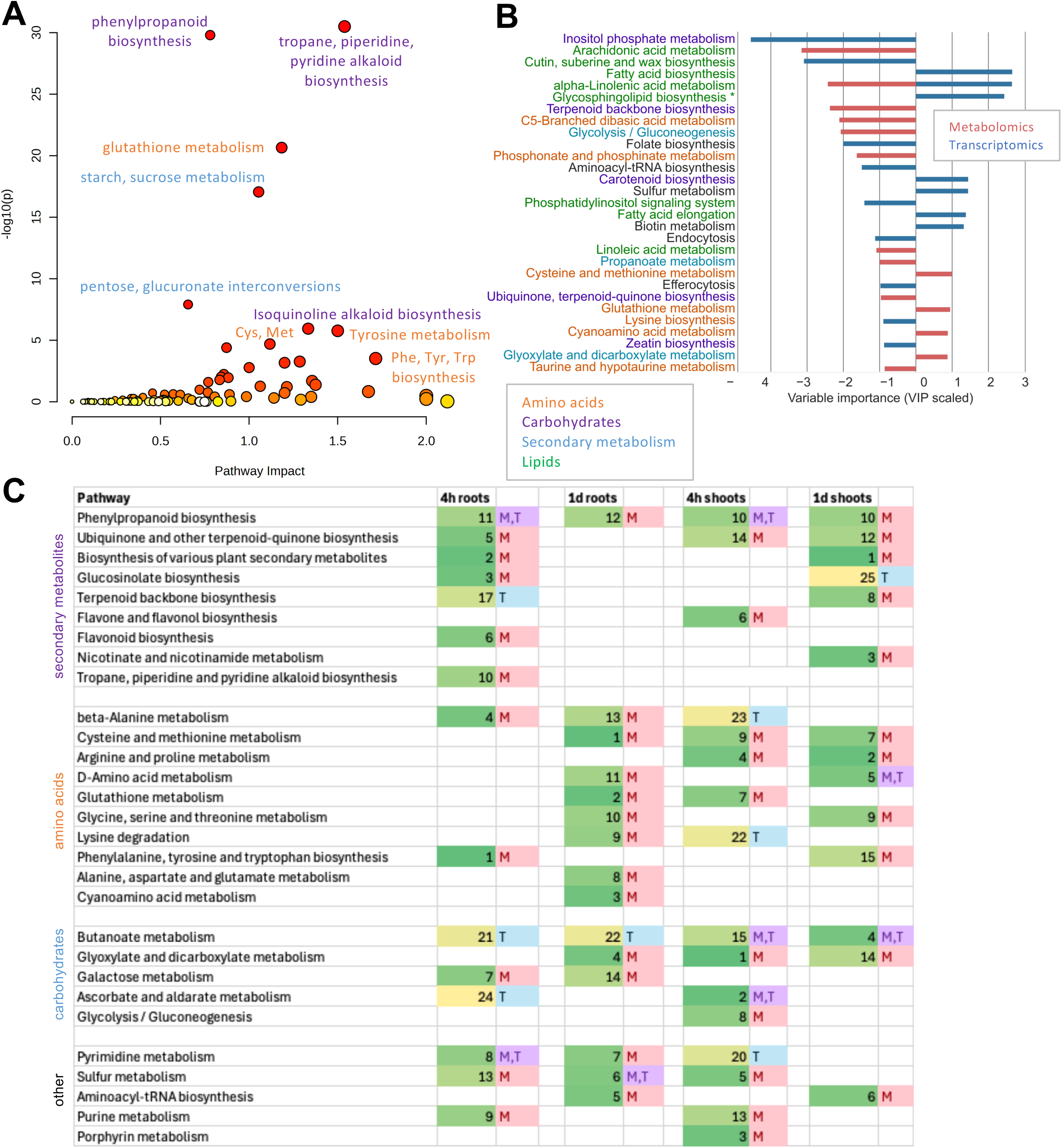
Multi-omics analysis highlights importance of amino acid metabolism. Multi-omic analysis integrating Metabolomic and transciroptomic datasets of 4 h and 1 d. A, MetaboAnalyst joint analysis of genes significantly different between mock- and fep-treated whole plant RNAseq analysis (log fold change of fep/mock) compared with metabolite log2 fold changes (fep/mock) in roots at 1 d. Significantly distinct metabolite pathways are listed in Figure S9a, a closeup of the graphic is presented in Figure S9b with additional pathway labels. Comparison of 1 d RNAseq with 1 d shoot metabolite data is presented in Figure S9c, and 4h RNAseq with 4 h root metabolite data in Figure S9d. B,C Pathintegrate multiview analysis of significantly different genes between mock- and fep-treated whole plant RNAseq analysis compared with metabolite data of the respective time points. B, Top30 pathways changing in roots at 1 d, C, top pathways changing across all comparisons. Red bars/”M”: significance detected in metabolomics dataset, Blue bars/”T”: significance detected in transcriptomic dataset, purple “M,T”: significance detected in both datsets. Numbers in C signify the rank number of the pathway in the dataset (1: most significant). Orange: amino acid-related pathway, blue: defense-related pathway, purple: carbohydrate-related pathway, green: fatty acid-related pathway. N =5961 significant genes at 4 h, 4587 significant genes at 1 d, 480 metabolites.

In the second analysis with PathIntegrate Multiview^20^, the same trends were iden6fied overall (Figure 3B, C): defense pathways ranked important (phenylpropanoids, alkaloids, terpenes, flavonoids, glucosinolates), as well as carbohydrates (glyoxylate& dicarboxylate, butanoate, ascorbate, galactose, glycolysis& gluconeogenesis), and amino acids (Phe, Tyr, Trp, Cys& Met, Ala, Arg& Pro, Gly& Ser& Thr, Lys). The Multiview tool enables to distinguish the contribution of the different -omics datasets to the output generated. With this analysis, the importance of metabolite data becomes evident, as 44/60 (73%) highly ranked pathways were iden6fied with metabolomics only, whereas 13/60 (13%) were iden6fied with transcriptomics only and 3/60 (14%) with the combination of both datasets (Figure 3C).

### Exudation is altered dynamically in response to elicitors

To investigate exudate profile changes in response to elicitors, plants were grown in semi-hydroponic systems and treated with elicitors in three distinct procedures, as it was to date unclear which treatment elicited a defense response. The plant medium was treated for 2 h with elicitors once at the beginning of the timecourse, with reversion back to the standard growth medium (pulse) or with constant elicitor presence (constant), or plants were vacuum infiltrated with medium containing elicitors (infiltration). Mock-treated plants were used as control (Figure 4A). Exudates were collected 2 h – 3 d after treatment for 2 h each. Shoots and roots were collected at the end of the experiment at 3 d. The elicitor treatments did not impact root- or shoot weight significantly (Figure S8A). Overall, 177 compounds were detected in exudates, 2045 in roots, and 1921 in shoots (Table S1). 152 compounds (86% of exudate metabolites, 7-8% of tissue metabolites) were detected in all mock-treated samples and 1769 were detected in tissues only (87-92%, Figure 4B). Few compounds were specific for a sample type except 124 compounds only detected in roots (6%). Of the 177 exuded compounds, 47% were distinct in any pairwise comparison. At the earliest timepoint (4 h after start of the timecourse, 2 h elicitor treatment and 2 h exudate collection), the exudate profiles of the different treatment regimes clearly clustered apart from mock-treated plants (Figure 4C). Overlap of pulse and constant treatments at this timepoint was to be expected, as the treatment was the same up to this timepoint. At the last timepoint, exudates of pulse and infiltration treated plants overlapped with mock treated plants, whereas exudates of plants continuously treated with elicitors remained distinct, indicating that the pulse and infiltration treatment did not elicit a lasting metabolic response (Figure 4D). PCA analyses of 6 h – 2 d timepoints revealed a slow movement of initially distinct pulse and infiltration treatments back to mock treatment (Figure S8B). Tissue metabolite profiles did not reveal significant differences at the end of the experiment (Figure S8C). Overall, 69 compounds were significantly distinct (Anova/Tukey) between mock and other treatments in at least one timepoint (38% of exuded metabolites). Clustering of the compounds revealed six clusters consisting of compounds belonging to various chemical classes (Figure 4E). Cluster I compounds decreased abundance at later timepoints, cluster II compounds increased early, and cluster V compounds, the largest cluster, increased early in infiltrated and late in constant-treated exudates. Other clusters showed a mix of higher and lower abundance depending on timepoint and treatment. For example, the phenylpropanoid cinnamate was increased consistently at 4 h and 6 h, whereas fumarate was decreased especially at 2 d and 3 d (Figure 4F).

**Figure 4:**
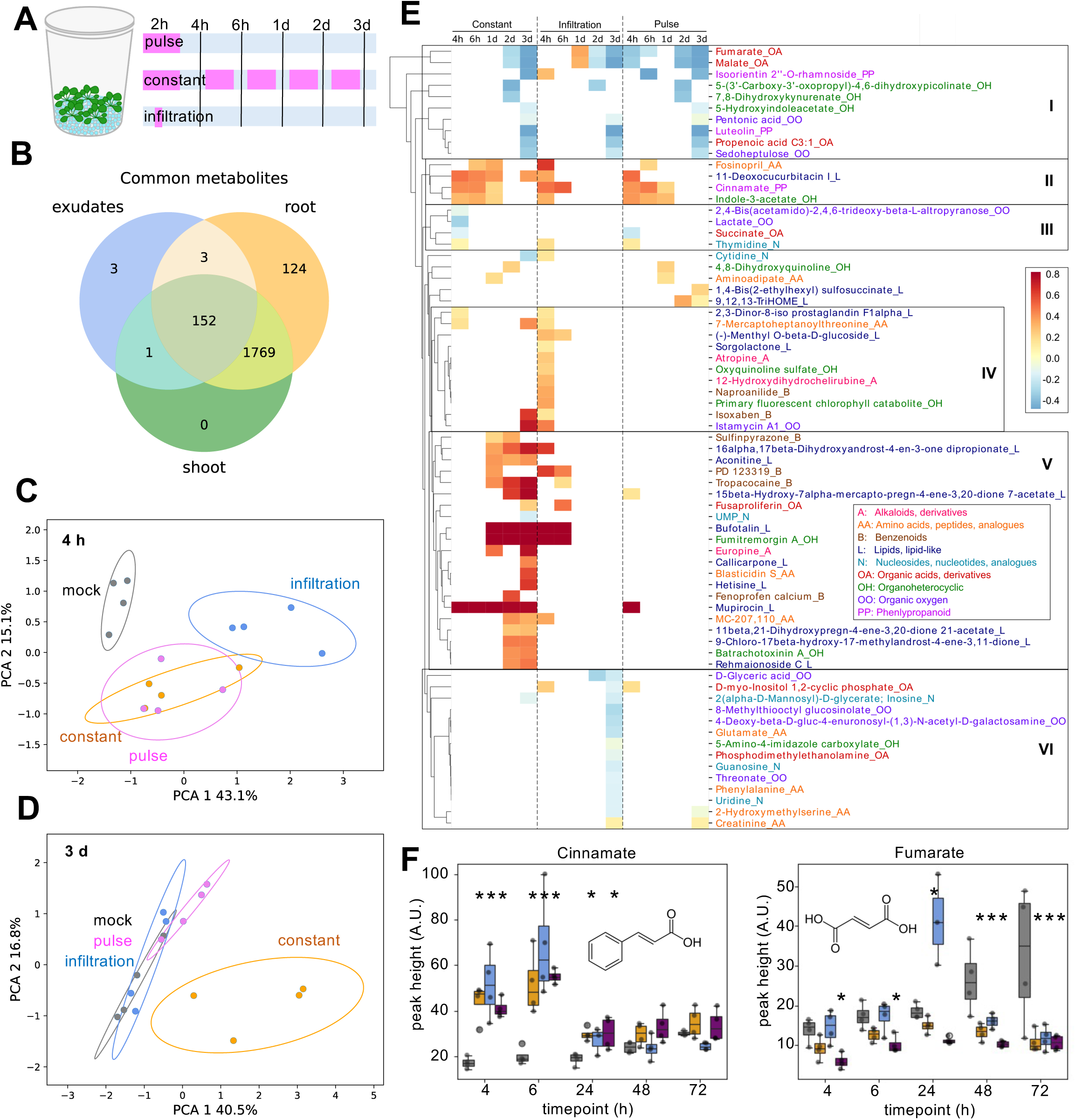
Elicitor treatment changes exudation dynamics. A, Experimental setup: 5 Arabidopsis thaliana plants were grown in a semi-hydroponic, sterile setup. The growth jars were assigned grouped randomly at the start of the experiment and treated with mock (0.5 MS growth medium), or an elicitor mix (1 µM flg22, 1 µM elf18, 5 µM pep1 in 0.5 MS) in pulse (2 h replacement of growth medium with elicitor mix), in constant (growth medium replacement with elicitor mix constantly between exudate collection timepoints), or infiltrated (2x 1 min vacuum application with plants submerged in elicitor mix). Exudates were collected for 2 h at indiated time points. B, Venn diagram of compounds present in shoot, root, exudates of mock-treated plants 3 d after beginning of the experiment. C-D, Principal component analysis (PCA) or exudate metabolite profiles at 4 h (C) or 3 d (D) after beginnig of the experiment. PC: Principal component with percent of variance explained. Other timepoints are displayed in Figure S8. E, clustered heatmap of fep/mock signal in phenylpropanoids for all experimental treatments. Color scale indicates higher than mock (red), lower than mock (blue), and not significantly changing (white). Six metabolite clusters were manually assigned, and compounds were colored according to Classyfire class. F, examples of compounds significantly changing between mock and elicitor-treated plants. A.U.: arbitrary units. Grey: mock, orange: constant, blue: infiltration, purple: pulse treatment. N = 4 jars with 5 plants each. Number of metabolites: 177 in exudates, 2045 in roots, 1921 in shoots.

### An integrated view on central carbon metabolism, amino acids, and defense

We displayed the transcriptomic and metabolomic data on a custom pathway map (Figure 5). Starting with carbon fixation and the citrate (TCA) cycle, few genes were differentially regulated (see also Figure S9), and the few metabolites detected in the dataset were less abundant in elicitor-treated samples compared to mock. Interestingly, carbohydrates were the only class showing a strong decrease of 20% of compounds in the entire dataset (Figure 1), likely correlating with the decrease of carbon fixation, ultimately resulting in the growth arrest of elicitor-treated plants (Figure S1).

**Figure 5:**
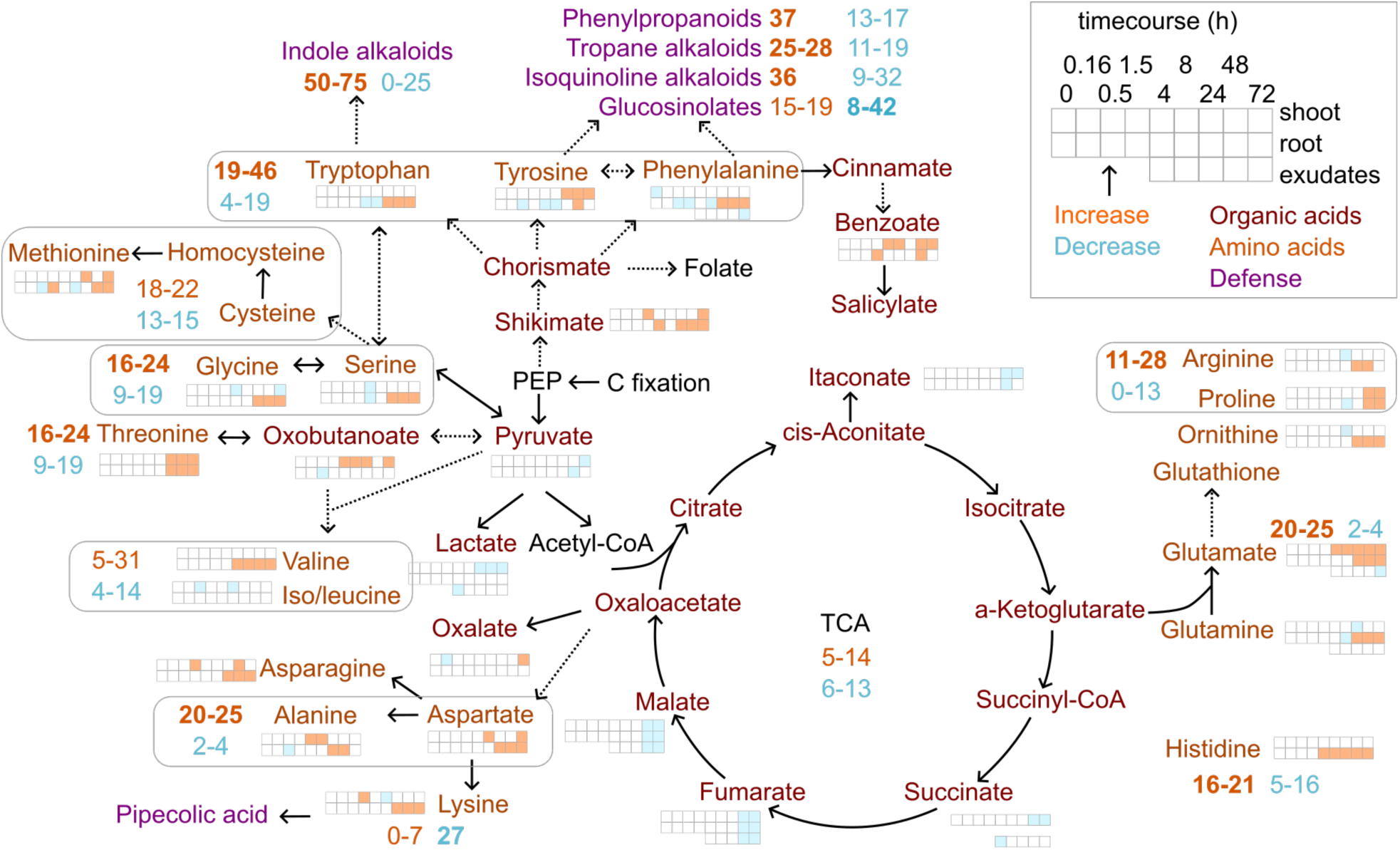
Integrated view on carbohydrate, amino acid, and defense metabolic pathways. Manual display of metabolites altered in various pathways. TCA: citric acid cycle. Bold lines: direct metabolite conversion, dashed lines: multiple steps. Metabolite colors: dark red: organic acids, orange: amino acids, purple: defense-related compounds. Significant increases (orange) and decreases (blue) for metabolite abundance in shoots (top line), roots (middle line), and exudates (bottom line) are indicated. The different timepoints are represented by squares. Missing squares: metabolite was not detected in dataset. Gene expression for different pathways are indicated in numbers as percent of genes with increased (orange) and decreased (blue) expression (see Figure S9). Bold numbers signify more genes changing.

For most amino acids, increased abundance was detected at later timepoints, sometimes accompanied by lower abundance at earlier timepoints (Figure 1G, Figure S5, Figure 5). Often, roots showed more significant changes than shoots, and many amino acids were absent from exudates (Figure S5, Figure 4, Figure 5). Genes assigned to specific amino acid pathways were more often up-than downregulated, although the transcriptional upregulation was less evident when compared to the metabolite data. Interestingly, the aromatic amino acids tryptophan, tyrosine, and phenylalanine that are precursors of many defense compounds ^1,2^ were not stronger regulated than other amino acids.

Defense metabolites increased in abundance upon elicitor treatment: Phenylpropanoids and isoquinoline alkaloids increased in roots especially at later timepoints (Figure 1). Multiple GO terms related to defense similarly increased in expression in the transcriptional data (Figure 2), and the integrated analysis confirmed the importance of phenylpropanoids, alkaloids, and other defense pathways (Figure 3).

Tryptophan as an indole alkaloid precursor increased in abundance in roots, as did expression of genes involved in tryptophan, tyrosine, and phenylalanine synthesis (Figure 5). Camalexin, the only indole alkaloid found in Arabidopsis^21^, was below detection limit with the metabolomics pipeline used here. Tyrosine and Phenylalanine are precursors for isoquinoline and tropane alkaloids, phenylpropanoids, and glucosinolates. They increased in expression and in abundance at later timepoints, preceded by a decrease in abundance at earlier timepoints (Figure 5). Genes associated with these defense pathways were mostly upregulated, but glucosinolate-related genes exhibited a decrease in expression (Figure 5). Phenylpropanoids increased first in roots followed by shoots at an early timepoint, followed by a general increase at later timepoints (Figure 1).

Apart from the aromatic amino acids, aliphatic amino acids also changed in abundance, and few of them were reported to be involved in biotic interactions. Glutamate for example is involved in pathogen and herbivore response in salicylic acid dependent and independent ways^22^. It increased in abundance in roots and shoots in later timepoints, whereas exudate levels remained unchanged (Figure 5). Also, expression of genes involved in this metabolic pathway increased (Figure 5). Lysine is a precursor for pipecolic acid synthesis, a long-distance signaling molecule involved in defense^23,24^. Lysine and methyl-lysine increased in roots in later timepoints and remained unchanged in shoots (Figure S5, Figure 5), and pipecolic acid and its derivatives were not detected in the metabolite dataset. Interestingly, gene expression of lysine synthesis is rather downregulated (Figure 5) and genes involved in pipecolic acid synthesis did not show a clear trend for altered expression (ALD1: decreased at 1 d, SARD4: increased at 1 d, UGT76B1: increased at 4 h)^25,26^. Further, the branched amino acids leucine, isoleucine, and valine were recently reported to suppress virulence of *Pseudomonas syringae*, suppressing host colonization, without affecting pathogen growth^27^. In our dataset, leucine and isoleucine could not be distinguished due to their shared molecular formula and were unchanged in response to elicitor treatment. Valine however was more abundant in roots, and could thus inhibit potential root-derived pathogens.

In summary, our data shows a decrease in carbohydrate- and TCA-cycle pathways early after elicitor treatment, and an increase of amino acids and defense-related compounds at later timepoints.

## Discussion

We find complex responses in tissues and exudates to elicitor treatment. Half of the compounds in shoots and roots changed in abundance, with approximately 10% at early (0- 30 min), 10-40% at middle (90 min-8 h), and 30-60% at late (1-3 d) timepoints. Elicitor-treated plants featured distinct exudation profiles at the earliest timepoint of collection (4 h), which lasted until 3 d when elicitors were present, and which diminished slowly over the course of 3 d when elicitors were only present for two hours at the start of the experiment. Responses in roots generally preceded shoots, which is explained with the experimental setup in which roots were treated with elicitors. The distribution of metabolite chemical classes did not change upon elicitor treatment. The different chemical classes showed various early-middle response pattern, but most featured a general increase in compound abundance at later time points. Defense compounds were upregulated (phenylpropanoids, alkaloids) in a pathway analysis, as were many amino acid-related pathways. The transcriptional analysis confirmed distinct responses at an early and late timepoint with early upregulation of immunity-related GO terms, and later upregulation of defense and few amino acid-related terms. The integrated metabolome-transcriptome analysis underlined the importance of the metabolite data, confirming differential regulation of defense-related and amino acid-related pathways. Interestingly, although mostly aromatic amino acids are well-recognized as precursors for defense compounds, we see an increased abundance for most amino acids.

Defense compounds can be grouped in three general classes, isoprenoids, shikimates, and alkaloids. All of these defense compounds are based on primary products of carbon fixation (pyruvate, glyceraldehyde-3-phsophate, acetyl-CoA). In our data, the TCA cycle is not strongly regulated, the few compounds detected in our dataset are downregulated at later timepoints, which might either indicate an inhibition of photosynthesis or increased use of TCA products (pyruvate, organic acids) for e.g. defense compound synthesis. Isoprenoids are synthesized via the MVA or MEP pathways to generate the C5-unit of terpenes^28^. MVA and MEP pathways are not well represented in our dataset, however, the shikimate pathway synthesizing aromatic acids and a suite of defense compounds is present. Shikimate, one of the core intermediates of the pathway, is upregulated mostly in roots at later timepoints, as are the aromatic amino acids. Up to 30% of fixed carbon is funneled into the shikimate pathway^29^: an upregulation of this pathway might thus explain the lower TCA metabolite levels. Tryptophan, one of the products of the shikimate pathway, then gives rise to tryptamine and indole alkaloids, which are upregulated in our transcriptional analysis. Tyrosine is converted to tyramine and isoquinoline alkaloids, which similarly exhibit transcriptional upregulation, and phenylalanine is a precursor to phenylpropanoids and other compounds, which are generally upregulated transcriptionally as well, with the exception of the glucosinolates that show a tendency for reduced expression ^1,2^.

Aside from the aromatic amino acids, many other amino acid levels were increased as well in our dataset. Many of these responses were more pronounced in roots than shoots. Elevated amino acid levels in response to pathogens were reported already in numerous studies investigating plants infected with bacteria, viruses, or fungi^3–5,22,30^. Compellingly, elevated amino acid levels were associated with susceptible ecotypes or cultivars, and generally not observed in resistant lines that feature higher levels of defense compounds. For example, in Arabidopsis ecotypes susceptible to *Pseudomonas syringae*, leucine and phenylalanine were enriched and sucrose levels decreased, whereas the resistant ecotype featured elevated glucosinolate levels ^5^. Another study found a consistent increase in glutamate and other amino acids in Arabidopsis ecotypes susceptible to a virus^4^. Interestingly, this study measured an increase in sucrose levels. Thus, the effect on central carbon metabolism seems to vary either with the pathogen or the timing of the assay but importantly, the increase in amino acid levels seems to be consistent. The metabolic changes we observe in our data are consistent with a susceptible line featuring enriched amino acids, decreased sugar and glucosinolate levels.

The importance of amino acids was somewhat highlighted in our transcriptomic dataset as well, with 1 GO-term being upregulated at 4 h, and 4 terms at 1 d after treatment. Specifically, the integrated multi-omics analysis helped to converge on essential pathways to focus our analysis: In our metabolomics dataset, many chemical classes increased abundance at later timepoints, and in our transcriptomic dataset, most GO-terms focused on biotic interaction terms. The integrated analysis then revealed four principal groups of regulated pathways, the first one being the defense compounds and the second one the amino acids (Figure 3). Multi-omic analysis is a relatively new strategy for plant biology research, but it has been instrumental to for example identify novel players in glucosinolate pathways by combining metabolomic and proteomic data^31^. Generally, combination of metabolomic with genetic analyses and bioassays is deemed an effcient avenue to identify breeding targets for biotic stress resistance^16^.

Our data and published reports indicate that elevated amino acid levels might be a key indicator of plant disease: a robust disease indicator should be present for a long time period in multiple tissues before the onset of visual symptoms. Determination of amino acid levels is likely a promising avenue for that: i) amino acids are quite abundant, making them detectable and quan6fiable with a variety of assays. ii) increases in many amino acid levels were detected as early as 8 h after elicitor treatment until the end of the experiment at 3 d. iii) In many studies, amino acid increases were detected 1-3 d after infection before the onset of visual symptoms^5,30^. iv) we detect amino acid increases across tissues, but more prominently in roots. It remains to be determined if the signal is stronger in leaves for leaf pathogens, and in roots for root pathogens. To turn this promising avenue into a tool, systematic studies should investigate whether amino acid increases are consistent responses to infection with various pathogens, for how long this increase persist, and which amino acids are the most reliable indicators (this might depend on the pathogen and host species used).

Amino acid levels could be elevated due to increased demand for defense compound synthesis, or due to a backlog caused for example by the observed growth arrest. Combining our data with published information, we can formulate the hypothesis that in pathogen-resistant lines, amino acids are depleted as building blocks for defense compounds and in susceptible lines, they might accumulate due to the growth arrest and lack of defense compound synthesis. A more detailed analysis of amino acid levels in response to different pathogens might aid in testing this hypothesis: Threonine, serine, arginine, and y-aminobutyric acid were recently reported as functional amino acids, impacting growth and development^32^. In our dataset, threonine increased in both tissues, arginine and serine in roots. This might provide a link to the growth arrest caused. In addition, pulse-chase experiments with labeled CO_2_ could aid to determine fluxes from central carbon metabolism to amino acids and defense compounds, determining altered fluxes in response to elicitors.

Additional experiments and technical advances would help to characterize the interplay of primary and secondary metabolism in defense responses further: the experiments presented here were performed with a flow injection metabolomics technique, without use of a chromatographic method. Use of polar and apolar chromatographies would aid in identifying compounds with higher reliability and broaden the number of compounds detected. In addition, for software-based analysis of metabolomic data, coverage of plant-specific compounds should be increased: in our dataset, only about half of compounds were part of the MetaboAnalyst database (Table S1). Especially the vast diversity of secondary metabolites detected in plants is missing in these databases. For example, the iPath3 software designed to map metabolites on KEGG pathways, lacks most of the secondary metabolite pathways (see for example Figure S4B). Despite these limitations, the experiments performed here point towards amino acids as central players in pathogen defense.

## Material& Methods

### Plant growth conditions

*Arabidopsis thaliana* Col-0 seeds were sterilized 15 min in 70% v/v ethanol followed by 15 min in 100% v/v ethanol and germinated on 0.5x Murashige and Skoog (MS) medium supplied with 0.7% w/v phytoagar, pH 5.7. Plates were placed vertically in a growth chamber with a 16 h light/8 h dark regime, 22°C at day and 18°C at night with an illumination of 150 μmol m^-2^ s^-1^. At 7 days after germination (dag), five plants were placed on square 0.5x MS plates with a width of 12 cm.

### Experimental setups

For the elicitor tissue timecourse, 50 ul of 1 uM flg22 (QRLSTGSRINSAKDDAAGLQIA), 1 uM elf18 (Ac-SKEKFERTKPHVNVGTIG), and 5 uM pep1 (ATKVKAKQRGKEKVSSGRPGQ) in 0.5x MS, or 0.5x MS as negative control, was added onto the root system of each plant at 18 dag. All peptides were synthesized by Scilight Biotechnology LLC (China) at 95% purity and dissolved in water. Shoot weights were determined, and roots and shoots were frozen separately in liquid nitrogen for metabolite extraction.

For the elicitor exudate timecourse, plants were transferred to a semi-hydroponic setup at 18 dag (McLaughlin Jove, xxx). At 21 dag, plants were treated as follows: for the pulse treatment, the 0.5x MS growth medium was replaced with 0.5x MS supplied with 1 uM flg22, 1 uM elf18, and 5 uM pep1 for 2 h, followed by the first exudate collection. For the constant treatment, the growth medium was similarly replaced with the elicitor mix for 2 h followed by exudate collection. In contrast to the other experimental treatments, the 0.5x MS growth medium was continuously supplied with elicitor mix. For the infiltration treatment, plants in jars were flooded with 0.5x MS supplied with elicitor mix, and infiltrated with a vacuum pump 2x 1 min. The medium was replaced with 0.5x MS without elicitors, and exudates were collected starting 2 h after start of the experiment in parallel with the other treatments. For the mock treatment, 0.5x MS without elicitors was used.

For exudate collection, the 0.5x MS growth medium was replaced with 20 mM sterile-filtered ammonium acetate, pH 5.7 for 2 h, and 2 ml of exudates were frozen for metabolite analysis.

### Sample preparation for metabolomics

Root and shoot tissues were ground at 1500 rpm for 1 min and extracted in 40:40:20 Methanol: Acetonitrile: ultrapure water twice for 1 h at 4°C with intermittent vortexing. Supernatants were pooled, dried under vacuum, and dissolved at a concentration of 70 mg 200 ul^-1^ in 1:1 MeOH: ultrapure water. Samples were diluted 1:150 before metabolomic analysis^33^. Root exudate volumes were adjusted according to root weight with 20 mM ammonium acetate before metabolomic analysis^19,33^.

### Metabolomics analysis pipeline

MeOH and ammonium acetate samples were prepared as blanks, and a standard mixture of 1 uM amino acids were used as quality control (QC) samples injected at the beginning and end of each 96-well plate to ensure consistent peak heights throughout the run. Samples were injected twice, and after each sample group (QC, exudates, tissues), blank samples were injected to reduce the probability of sample carryover.

A flow injection – time of flight mass spectrometry system of an isocratic Agilent 1200 pump with a Gerstel MPS2 autosampler and an Agilent 6550 QTOF mass spectrometer (Agilent, Santa Clara, CA, USA) was used for the experiments, with sewngs as published^34^. In brief, the mobile phase was 60:40% v/v isopropanol: ultrapure water supplied with 5 mM ammonium fluoride (negative ionization mode) with a flow rate of 150 µl min^-1^. Homoserine and Hexakis (1H, 1H, 3H-tetrafluoropropoxy)-phosphazine were used for mass correction. Mass spectra were recorded from 50-1000 m/z with a frequency of 1.4 spectra s^-1^. Source temperature was 325°C, with 5 ml min^-1^ drying gas and a nebulizer pressure of 30 spig. Fragmentor, skimmer, and octupole voltages were set to 175 V, 65 V, and 750 V. Spectral data was processed as published ^34^ and annotated based on the Kyoto Encyclopedia of Genes and Genomes (KEGG) and Human Metabolome Database (HMDB) databases.

Metabolite data was first compared against experimental blanks: a metabolite was kept if its peak height in any sample group (any timepoint and treatment) was 2x higher than in the experimental blank sample. Significantly distinct metabolites were determined with an Anova / Tukey post-hoc test comparing mock-versus elicitor-treated samples. Data were visualized with Python custom code for principal component analyses and hierarchical clustering.

Chemical classes were assigned with ClassyFire^35^. For this, the first Kegg ID was converted to international chemical iden6fiers InChI, which was used to query the ClassyFire tool (https://cx.fiehnlab.ucdavis.edu/).

Data was visualized with MetaboAnalyst 6.0 (Pang 024 NAR). Fold changes of elicitor treated / mock metabolite abundances were used as input for the Pathway Analysis tool (https://www.metaboanalyst.ca/MetaboAnalyst/ModuleView.xhtml). The pathways were manually labeled for visualization. Metabolites assigned to specific pathways were in addition mapped to the KEGG pathway maps with iPATH3.0^36^.

### RNA extraction and RNASeq

Plants were grown in the semi-hydroponic setup. At 20 dag, they were treated with 0.5x MS supplied with 1 uM flg22, 1 uM elf18, and 5 uM pep1 for 2 h, followed by incubation in 0.5x MS. Four hours or 1 d after treatment start, 50 mg of root and shoot tissues were sampled from 3-4 individuals and immediately frozen in liquid nitrogen. RNA was extracted according to manufacturers protocol (RNeasy Plant Mini kit, Qiagen, USA). The Functional Genomics Center Zurich prepared the RNAseq libraries and performed the sequencing: The quality of the isolated RNA was determined with a Qubit® (1.0) Fluorometer (Life Technologies, California, USA) and a Fragment Analyzer (Agilent, Santa Clara, California, USA). Only those samples with a 260 nm/280 nm ratio between 1.8–2.1 and a 28S/18S ratio within 1.5–2 were further processed. The TruSeq Stranded mRNA (Illumina, Inc, California, USA) was used in the succeeding steps. Briefly, total RNA samples (100-1000 ng) were polyA enriched and then reverse-transcribed into double-stranded cDNA. The cDNA samples was fragmented, end-repaired and adenylated before ligation of TruSeq adapters containing unique dual indices (UDI) for multiplexing. Fragments containing TruSeq adapters on both ends were selectively enriched with PCR. The quality and quantity of the enriched libraries were validated using Qubit® (1.0) Fluorometer and the Fragment Analyzer (Agilent, Santa Clara, California, USA). The product is a smear with an average fragment size of approximately 260 bp. The libraries were normalized to 10nM in Tris-Cl 10 mM, pH8.5 with 0.1% v/v Tween20. The NovaseqX (Illumina, Inc, California, USA) was used for cluster generation and sequencing according to standard protocol. Sequencing were paired end at 2 x150 bp.

Analysis of the read data was performed using the SUSHI framework^37^. Reads were preproscessed with fastp to trim adapters^38^. Read alignment was done using the STAR aligner^39^ with the TAIR10 reference. Gene-level expression values were generated using the ‘featureCounts’ function of the R package Rsubread^40^.

Transcripts per kilobase million (tpm) data was used for a principal component analysis. The non-normalized sequence read counts were processed with the Python DESeq2 workflow (https://github.com/owkin/PyDESeq2), a python implementation of the DESeq2 pipeline for bulk RNAseq for differential expression analysis^41,42^.

### Multi-omic data integration

RNASeq and metabolomics data was integrated with the PathIntegrate tool (https://github.com/cwieder/PathIntegrate). Briefly, genes with significantly different expression levels for mock versus elicitor-treated plants harvested 4 h or 1 d after treatment as defined by the DESeq2 workflow were used as input together with root or shoot metabolite data collected 4 h or 1 d after elicitor treatment. Arabidopsis metabolite pathways were downloaded from Reactome (https://reactome.org/), and the pipeline for multi-view analysis was followed^20^.

## Acknowledgements

We would like to thank Alexandra Siffert and Sarah McLaughlin for experimental support, Nicola Zamboni for generating metabolome data and Cecilia Wieder for support with implementing the PathIntegrate pipeline. At the functional genomics center, we would like to acknowledge Maria Domenica Moccia, Lennart Opitz, and Hubert Rehrauer for experimental suggestions, preparation of RNAseq libraries, and sequencing.

The work presented here was supported by the Swiss National Science Foundation (PR00P3_185831) to J.S., and by a Plant Science Center-Syngenta research fellowship awarded to K.S. and J.S., supporting C.J.

---

**Supplemental Figure 1.** A, Shoot weight of 3-week-old A. thaliana Col-0 at the time of harvest for mock and fep-treated plants. Data = average +-S.E.M., n=5 biological replicates of each 4-5 plants. B, Principal component analysis plots of metabolites extracted from roots and shoots mock- and fep-treated, grouped by tissue and/or timepoint. Percent of variability explained by PC1 and 2 are indicated, n = 3-5 biological replicates per tissue and timepoint. Number of metabolites: 480.

**Supplemental Figure 2.** A, chemical classes (Classyfire super-/subclasses) of roots and shoots in absolute numbers and percent for compounds present and compounds distinct between mock and fep-treated samples at any of the timepoints. Classes displayed as ‘other’ in Figure 1B are indicated in grey font. B, percent of compounds distinct between fep and mock treated tissues for all timepoints for additional chemical classes. Solid line: upregulated metabolites, dotted line: downregulated metabolites, green: shoot metabolites, brown: root metabolites. Arrows highlight patterns observed (same color, pattern as for lines, black: observ ed in both). N: number of metabolites present in each chemical class. Additional plots are shown in Figure 1D-G.

**Supplemental Figure 3.** MetaboAnalyst Pathway analysis of fep/mock ratio. A, closeup of differentially regulated root pathways, closeup of plot in Figure 1H. B, Differentially regulated shoot pathways. Blue: amino acid-related pathways, green: defense-related pathways, purple: carbohydrate-related pathways.

**Supplemental Figure 4.** KEGG maps with selected amino acids (A) and other pathways (B) distinct in MetaboAnalyst analyses of fep/mock root compounds (see also Fig 1H, Supp Fig 3). Bold: compounds higher abundant, italics: compounds lower abundant in fep-treated roots at significantly different timepoints compared to mock-treated roots. Regular font: compounds with complex lower/higher abundant responses across timepoints. Maps were built with iPath3.

**Supplemental Figure 5.** Clustered heatmaps of compounds significantly distinct between fep- and mock-treated tissues. Data averaged by timepoint, log transformed and centered around 0 (red: higher, blue: lower abundant, white: not significantly different from mock). A, shoot phenylpropanoids colored by chemical class (root phenylpropanoids displayed in Figure 1I). B,C, amino acids distinct in roots (B) and shooots (C).

**Supplemental Figure 6.** Principal component analysis of RNAseq data as transcripts per kilobase million reads. Percent of variability explained by PC1 and 2 are indicated, n = 4 biological replicates.

**Supplemental Figure 7.** A, Top pathways of MetaboAnalyst integrated analysis of roots at 1 d (for corresponding plot, see Figure 3a). The second column indicates how many genes in the pathways are matched, and the third column expressed the percent match, colored by green (highest) to yellow (lowest). B-D, MetaboAnalyst integrated analysis of genes significantly different between mock- and fep-treated whole plant RNAseq analysis compared with metabolite fold changes (fep/mock) of shoots at 1d (B), of roots at 4 h (C), and of shoots at 4 h (D). pathways in bold are also present in roots 1 d analysis, see Figure 3a. Orange: amino acid-related pathway, blue: defense-related pathway, purple: carbohydrate-related pathway, green: fatty acid-related pathway. N =5961 significant genes at 4 h, 4587 significant genes at 1 d, 480 metabolites.

**Supplemental Figure 8:** A, Root and shoot weight of plants in jars 3 d after experiment start. No significant differences are detected between mock and elicitor treatments. B, Principal component analysis of exudate metabolite profiles 6 h, 1 d, and 2 d after treatment. C, root and shoot metabolite profiles of plants 3 d after experiment start. Additional PCA plots are displayed in Figure 4. PC: Principal component with percent of variance explained.N = 4 jars with 5 plants each. Number of metabolites: 177 in exudates, 2045 in roots, 1921 in shoots.

**Supplemental Figure 9:** Selected Reactome pathways (black: TCA, orange: amino acid, blue: defense, black: plant pathogen) with number of genes associated with the pathway and percent of genes up- and downregulated in the transcriptomic dataset. Color code indicates low (red), medium (yellow), and high (green) percentage of genes changing.

